# A Comparative Analysis of Novel Deep Learning and Ensemble Learning Models to Predict the Allergenicity of Food Proteins

**DOI:** 10.1101/2021.03.10.434710

**Authors:** Liyang Wang, Dantong Niu, Xinjie Zhao, Xiaoya Wang, Mengzhen Hao, Huilian Che

## Abstract

Traditional food allergen identification mainly relies on in vivo and in vitro experiments, which often needs a long period and high cost. The artificial intelligence (AI)-driven rapid food allergen identification method has solved the above mentioned two drawbacks and is becoming an efficient auxiliary tool. Aiming to overcome the limitations of lower accuracy of traditional machine learning models in predicting the allergenicity of food proteins, this work proposed to introduce deep learning model - transformer with self-attention mechanism, ensemble learning models (representative as Light Gradient Boosting Machine (LightGBM) eXtreme Gradient Boosting (XGBoost)) to solve the problem. In order to highlight the superiority of the proposed novel method, the study also selected various commonly used machine learning models as the baseline classifiers. The results of 5-fold cross-validation showed that the AUC of the deep model was the highest (0.9578), which was better than the ensemble learning and baseline algorithms. But the deep model need to be pre-trained, and the training cost is the highest. By comparing the characteristics of the transformer model and boosting models, it can be analyzed that, each model has its own advantage, which provides novel clues and inspiration for the rapid prediction of food allergens in the future.

## 1. Introduction

Food allergy refers to inflammation of the human body caused by the body’s specific immune response through ingestion, inhalation, or skin contact with specific types of food proteins. It belongs to ones of allergic diseases. In recent years, people’s attention to food allergy has been increasing because it will cause a series of complications [1]. For example, the most common manifestations of extra-intestinal symptoms of food allergies are angioedema, various skin rashes, and eczema. It can also cause rhinitis, conjunctivitis, recurrent oral ulcers, bronchial asthma, allergic purpura, arrhythmia, headache, and dizziness, and even lead to systemic reactions of anaphylactic shock. Meanwhile, the increasing prevalence of food allergies and the significant positive correlation between food allergies and the respiratory tract are becoming one of the main problems threatening human health [2,3]. Studies reveal that the occurrence rate of respiratory diseases in patients with food allergies is significantly higher than that in patients without food allergies [4]. Food allergies are mainly induced by food allergens, which are food antigen molecules that can cause immune responses to the human body. Almost all food allergens are proteins, most of which are water-soluble glycoproteins with a relative molecular mass between 10,000 and 70,000.

According to the report by the United Nations Food and Agriculture Organization, more than 90% of the food allergens have existed in eight types of food: fish, eggs, milk, crustaceans, soybeans, peanuts, nuts, and wheat [5]. Although more than 180 kinds of foods have been identified as containing allergens, there are still many food allergens that have not been discovered. Therefore, the identification of food allergens are particularly important. Traditional food allergen identification methods can be divided into two categories: serology and cytology. The serological method judges whether a protein is an allergen based on the binding ability of the test protein to the IgE in the positive serum. The cytological method is based on the inflammatory mediators produced by the immune cells when it is stimulated by the test protein to evaluate its sensitization. In addition, based on the different locations where the detection method is performed, the allergy identification method can also be divided into in vivo methods and in vitro methods. In vivo tests include double-blind placebo-controlled food challenge test (DBPCFC), skin test (ST), and animal models; in vitro tests include enzyme-linked immunosorbent assay (ELISA), simulated gastrointestinal digestion, histamine release test, western blotting, and Allergen adsorption experiment. There is no doubt that the results of the above traditional identification methods are relatively reliable. However, they have the disadvantages of a long experimental period and high cost, leading to difficulties in high-throughput and high-speed food allergen prediction.

At present, bioinformatics methods have been widely used in food allergen detection. The most typical one is the basic local alignment search tool (BLAST), BLAST can discover and screen out similar sequences by comparing the tested sequences with various nucleotide sequences and protein sequences in the National Center for Biotechnology Information (NCBI) database, which can be used to predict and analyze the functional structure of proteins and nucleic acids. In [6], the author used BLAST to identify potential food allergy cross-reactions and achieved the desired results. Goodman [7] et.al employed bioinformatics tools including BLAST to identify allergens, proteins that are very similar to allergens, and the allergic cross-reactions they may induce. It should be emphasized that although BLAST has a higher efficiency in comparison of sequences and is relatively convenient to conduct, there is a greater probability of false positives in the comparison results [8].

With the development of artificial intelligence technology, machine learning has been gradually applied in the prediction of protein functions, and good results have been obtained. Using a labeled protein database, researchers train neural networks to study the primary or secondary structure of the protein, discover the in-depth relationship between the sequence structure of physical and chemical properties of the proteins and their functions. Then the well-trained neural network can be used to predict unknown proteins. In the early years of research on allergenicity prediction, Soeria Atmadja [9] et.al performed supervised learning on the extracted peptide sequence features and selected support vector machines (SVM) with linear kernel functions for classification to obtain relatively accurate results. In [10], classifiers, such as K-nearest neighbor (K-NN), were adopted to predict the allergenicity of allergens. Besides, some researchers have extracted the pseudo-amino acid composition (PseAAC) feature of the allergen and employed the SVM classifier to predict the protein’s allergenicity [11]. In recent years, deep learning models, such as deep neural networks (DNN), have also been used for the identification of allergens [12], which behave better than traditional machine learning methods. Meanwhile, online servers developed by using various machine learning algorithms are successively reported recently, which greatly improves the efficiency and facilitates high-throughput prediction [13–15]. The efficiency of the prediction method based on machine learning algorithms is much higher than in vivo and in vitro experiments. Furthermore, the accuracy of the predictions is constantly breaking through with the improvement and optimization of the model. DNNs have become the mainstream tool for food allergen prediction in the future.

Bidirectional Encoder Representation from Transformers (BERT) is mainly used for natural language processing (NLP) and currently is rarely applied for peptide or protein function prediction [16]. We found that it can extract high-dimensional features between peptides sequence for study, which is a novel prediction method. Convolutional Recurrent Neural Network that (CRNN) is used for end-to-end recognition of text sequences of indefinite length. Instead of cutting out a single text first, it converts text recognition into a sequence-dependent sequence learning problem. It has also been reported CRNN plays a role in the function prediction of proteins [17]. The novel ensemble learning model is becoming one of the mainstream methods to improve the performance of machine learning, which has shown superior performance compared to traditional classifiers in text classification [18], disease diagnosis [19], and other fields, and its application in the peptide sequence classification has been rarely seen. The three methods above all provide novel ideas to improve the accuracy and performance of machine learning algorithms to predict the allergenicity of food proteins.

In this paper, we originally introduced BERT, a novel pre-training model in the field of natural language processing, into the allergenicity prediction of food allergens. An independent attention mechanism in each layer was adopted, so compared to traditional Recurrent Neural Networks (RNN), our network can capture longer-distance dependencies more efficiently. Additionally, in order to make a comparison on characteristics between the deep learning model and ensemble learning model in this task, 2 novel ensemble learning models-LightGBM and XGBoost for 5-fold cross-validation were employed. The results showed that for the dataset in this work, the introduced ensemble learning models (LightGBM, XGBoost) were better than the baseline classifiers, but didn’t perform as well as deep learning. However, the convenience brought by its short training time makes it suitable for certain specific environments. The novel self-attention mechanism of BERT with the superior performance has infinite potential in larger-scale data training in the near future.

## 2. Materials and Methods

The whole method of this work is shown in Figure **1**.

**Figure 1.**
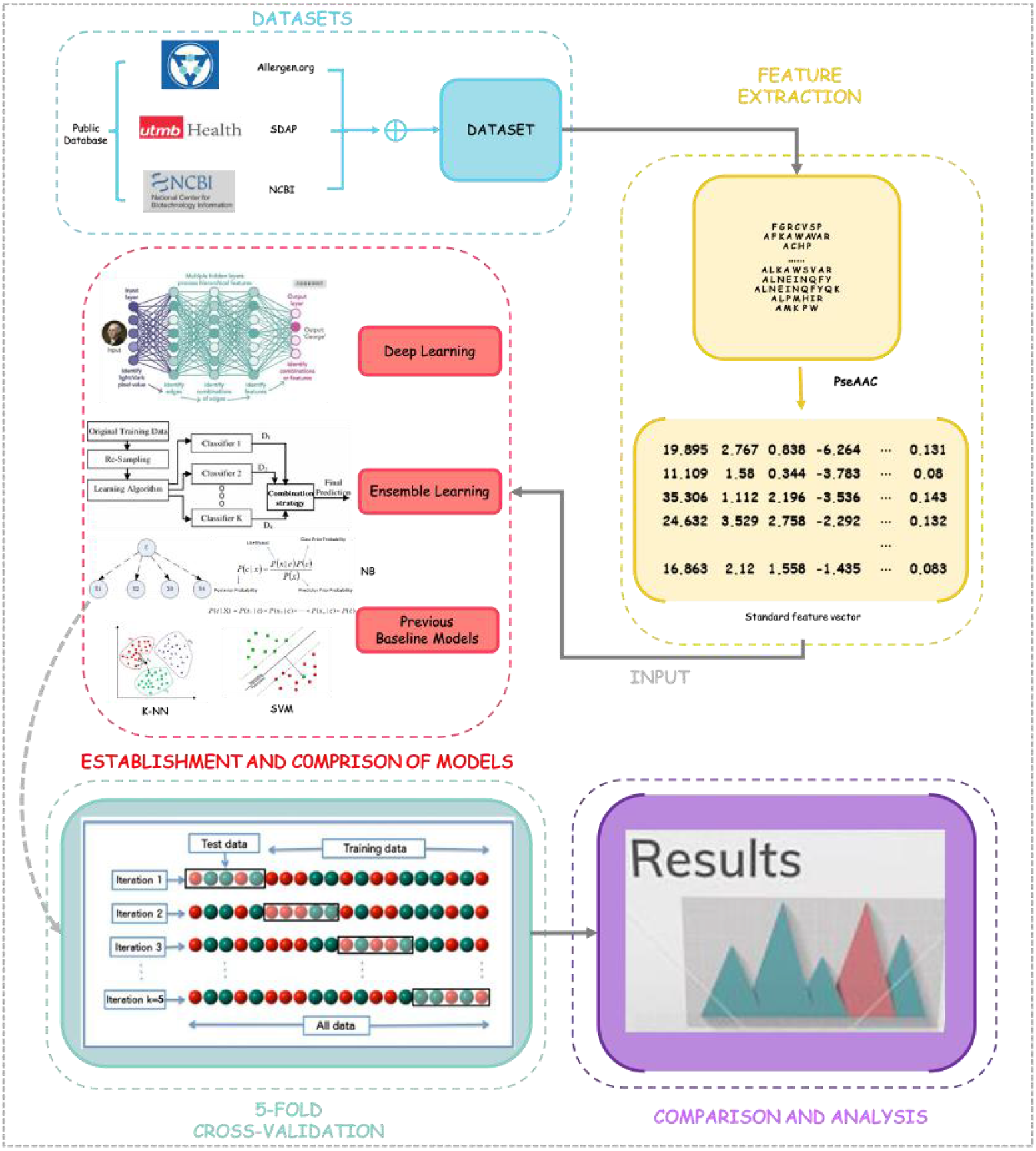
The workflow of this study.

### 2.1 Construction of Datasets

The food allergen datasets adopted in this study are from Allergen Nomenclature (http://www.allergen.org/index.php), Structural Database of Allergenic Proteins (SDAP) (http://fermi.utmb.edu/SDAP/), and NCBI (https://www.ncbi.nlm.nih.gov/) three public databases. This research has gathered 583 food allergens that were officially reported to be allergenic and corresponding protein sequences as positive samples, and 600 food proteins (not reported as allergens) and corresponding sequences as negative samples. The dataset has been rigorously screened, and there is no duplication between positive and negative samples.

### 2.2 Representation of Sequences of Food Allergens

Pseudo-amino acid composition (PseAAC) was first proposed by Chou [20], and it is one of the classic protein sequence feature representation methods. The type II PseAAC of a protein can be expressed as a 20+iλ-dimensional feature vector, where the first 20 dimensions reflect the frequency distribution of each amino acid on the protein, and i represents the number of amino acid properties used when generating PseAAC (hydrophilicity, hydrophobicity, etc.), λ represents the sequence correlation factor. Therefore, PseAAC simultaneously contains amino acid’ s composition and sequence information and the interaction information between them. In this research, we considered 6 properties (hydrophobicity, hydrophilicity, mass, pK1(a-CO2H), pK2(NH3) and pI(at 25°)), i was set to 6, λ was set to 5, and the weight factor ω was set to 0.05. As a result, the fixed dimension of the PseAAC feature vector of the input machine learning models (except for BERT, because it comes with a dictionary) was 50 dimensions.

### 2.3 Artificial Intelligence Models

This section mainly introduces the artificial intelligence models adopted in this work. Among them, the focus is on the deep model-BERT algorithm and the novel boosting model-LightGBM, which highlights their superior mechanism.

#### 2.3.1. Deep Learning Model

BERT is a self-supervised method for pre-training deep transformer encoders, which can be finetuned for different downstream tasks after pre-training. BERT can be optimized for two training objectives-mask language modeling (MLM) and next sentence prediction (NSP), and only large unlabeled datasets are needed for its training. As a novel deep learning model, BERT is commonly used in the field of NLP, and it is rarely applied in the study of food allergen prediction.

The architecture of BERT is a multi-layer transformer structure. Transformer is an encoder-decoder structure formed by stacking several encoders and decoders. The encoder consists of Multi-Head Attention and a feedforward neural network, which is used to convert the input protein sequence into a feature vector (Figure **2**). The input of the decoder is the output of the encoder and the predicted result, which is composed of Masked Multi-Head Attention and a feedforward neural network. The decoder outputs the conditional probability of the final result (Figure **2**). The highlight of BERT is the use of Multi-Head Attention, which divides a word vector into N dimensions. Since the allergen sequence is mapped in the high-dimensional space in the form of multi-dimension vectors, the mechanism of Multi-Head Attention enables the model to learn different characteristics of each dimension. The information learned from adjacent spaces is similar, which is more reasonable than mapping the entire space together.

**Figure 2.**
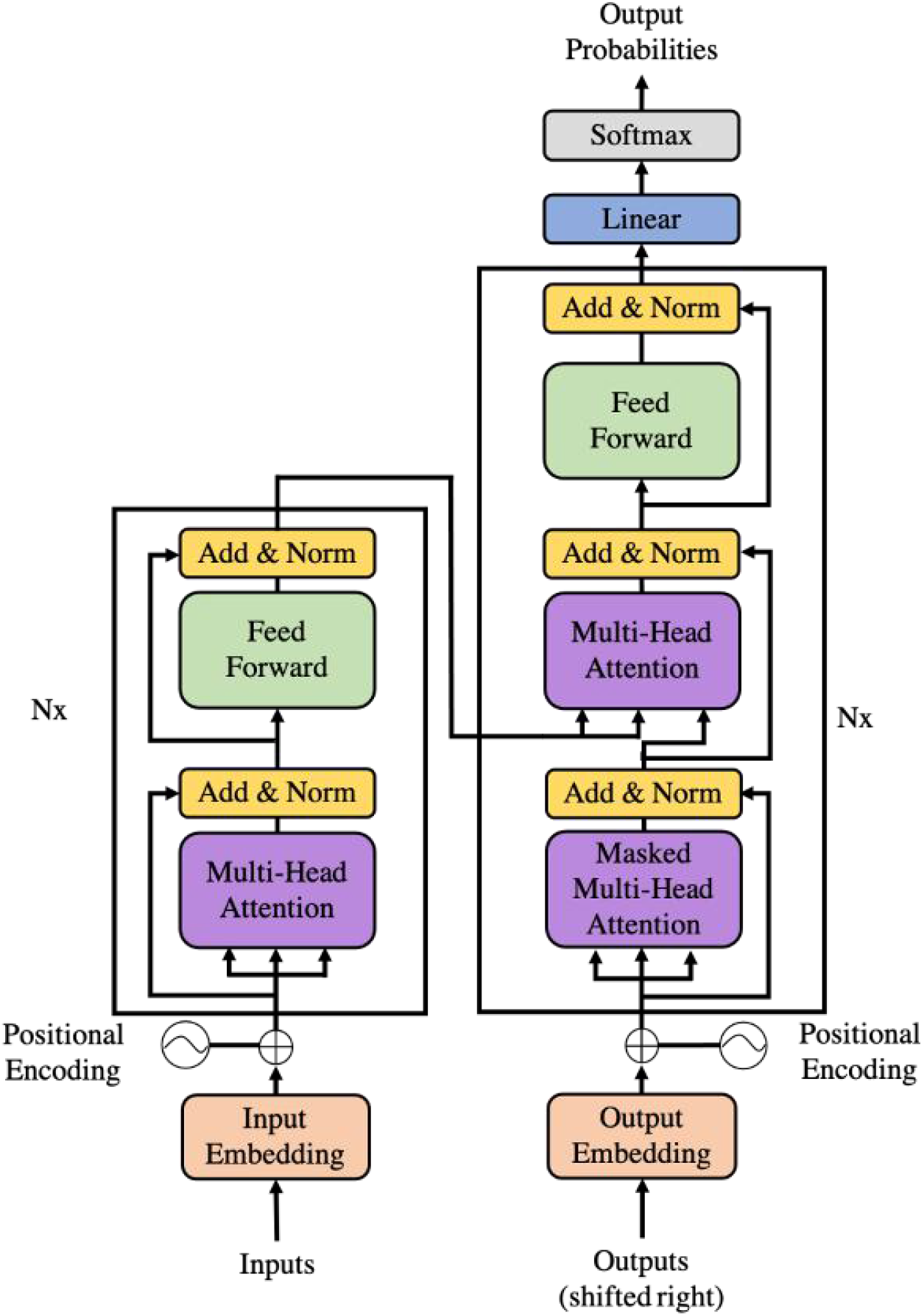
Transformer structure.

In this study, we employed the pre-training model, protBERT (specially trained from protein sequences) [21], which transferred a large number of operations deployed in specific downstream NLP tasks to pre-training word vectors. After obtaining the word vector used by BERT, a multi-layer perceptron (MLP) to the word vector was added. This experiment separated each amino acid character with a space and cut the amino acid sequence so that the amino acid chain formed a string with a certain length, which was used as a basic structure input.

#### 2.3.2. Ensemble Learning Models

##### 2.3.2.1 Light Gradient Boosting Machine (LightGBM)

LightGBM was proposed by Microsoft in 2017. It is a novel Gradient Boosting Decision Tree (GBDT) algorithm framework. It currently shows excellent results in economic forecasting, disease diagnosis and other fields [22,23], but little information about its application in food allergen predictions has been reported so far. In order to solve the time-consuming problem of traditional GBDT when the training dataset is large and complicated, LightGBM model further uses two methods and further improves the accuracy of the model.

One is GOSS (Gradient-based One-Side Sampling, gradient-based one-side sampling). Instead of using the sample points to calculate the gradient, GOSS calculates the gradient by sampling the samples. GOSS excludes most of the samples with small gradients, and only employs the remaining samples in the calculation. Although GBDT does not have data weights, each data instance has a different gradient. According to the calculated definition of information gain, instances with large gradients have a greater impact on information gain. Therefore, when downsampling, samples with large gradients should be kept as much as possible (screened with predefined threshold or highest percentiles), and samples with small gradients should be randomly removed. Experiments show that this measure can obtain more accurate results than random sampling with the same sampling rate, especially when the range of information gain is large.

The second is EFB (Exclusive Feature Bundling). Instead of using all features for scanning to obtain the best segmentation point, some features are bundled together to reduce the dimension of the feature. A Histogram algorithm is employed in LightGBM. The basic idea is to discretize continuous eigenvalues into k integers, and construct a histogram with the width of k. When traversing the data, the discretized value is used as the index to accumulate statistics in the histogram. After traversing the data once, the histogram accumulates the required statistics. Then according to the discrete value of the histogram, an optimal split point can be found by traversing the data again (Figure **3**). This mechanism reduces memory usage and speeds up model training.

**Figure 3.**
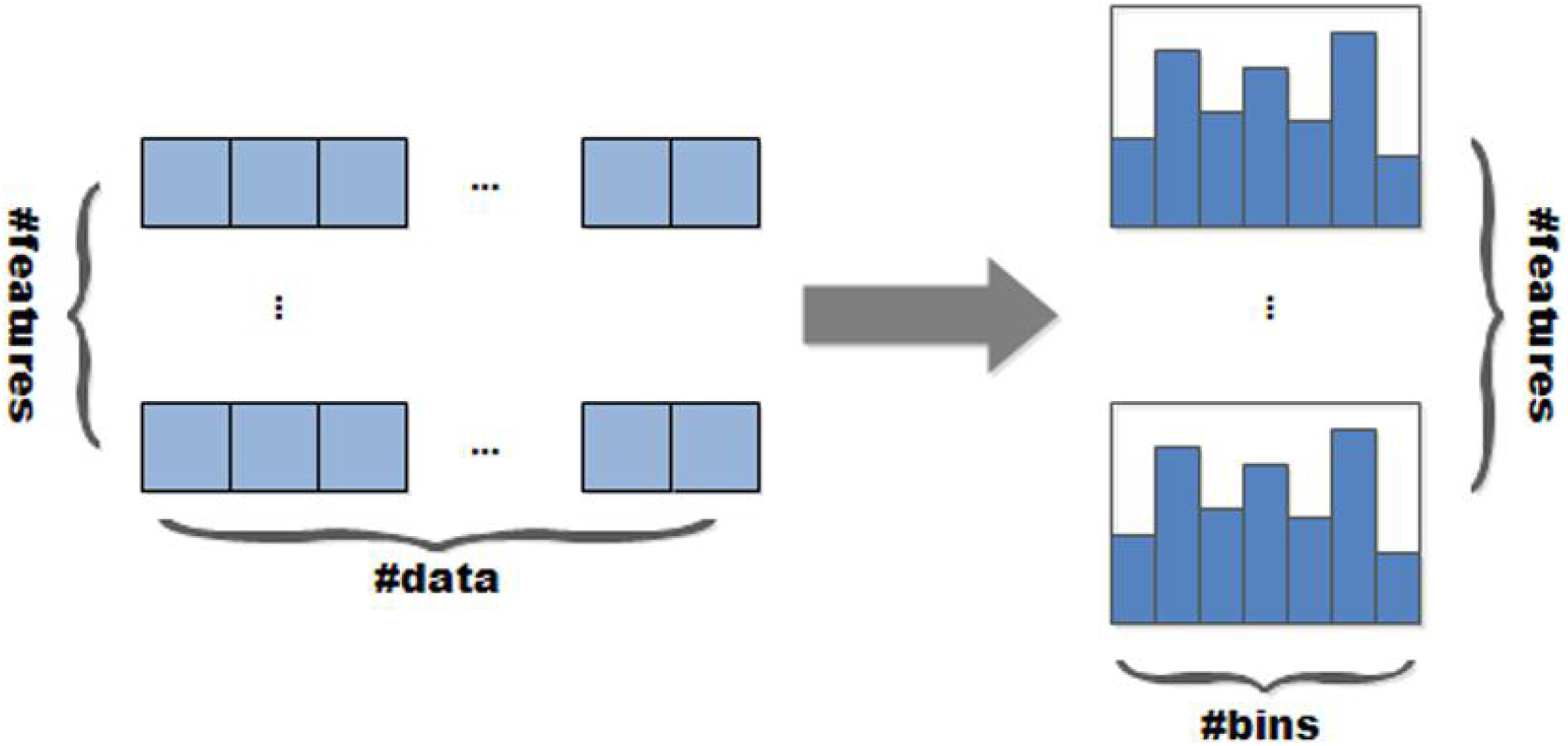
Histogram algorithm flow.

Furthermore, LightGBM adopts a Leaf-wise strategy to construct the tree models. Each time, the leaf with the largest split gain in all current leaves is chosen to split and the process is repeated. Compared with the traditional Level-wise strategy, this strategy can reduce more errors and get better accuracy with the same number splits. Meanwhile, the parameter max_depth is introduced to limit the depth of the tree and avoid overfitting as shown in Figure **4**.

**Figure 4.**
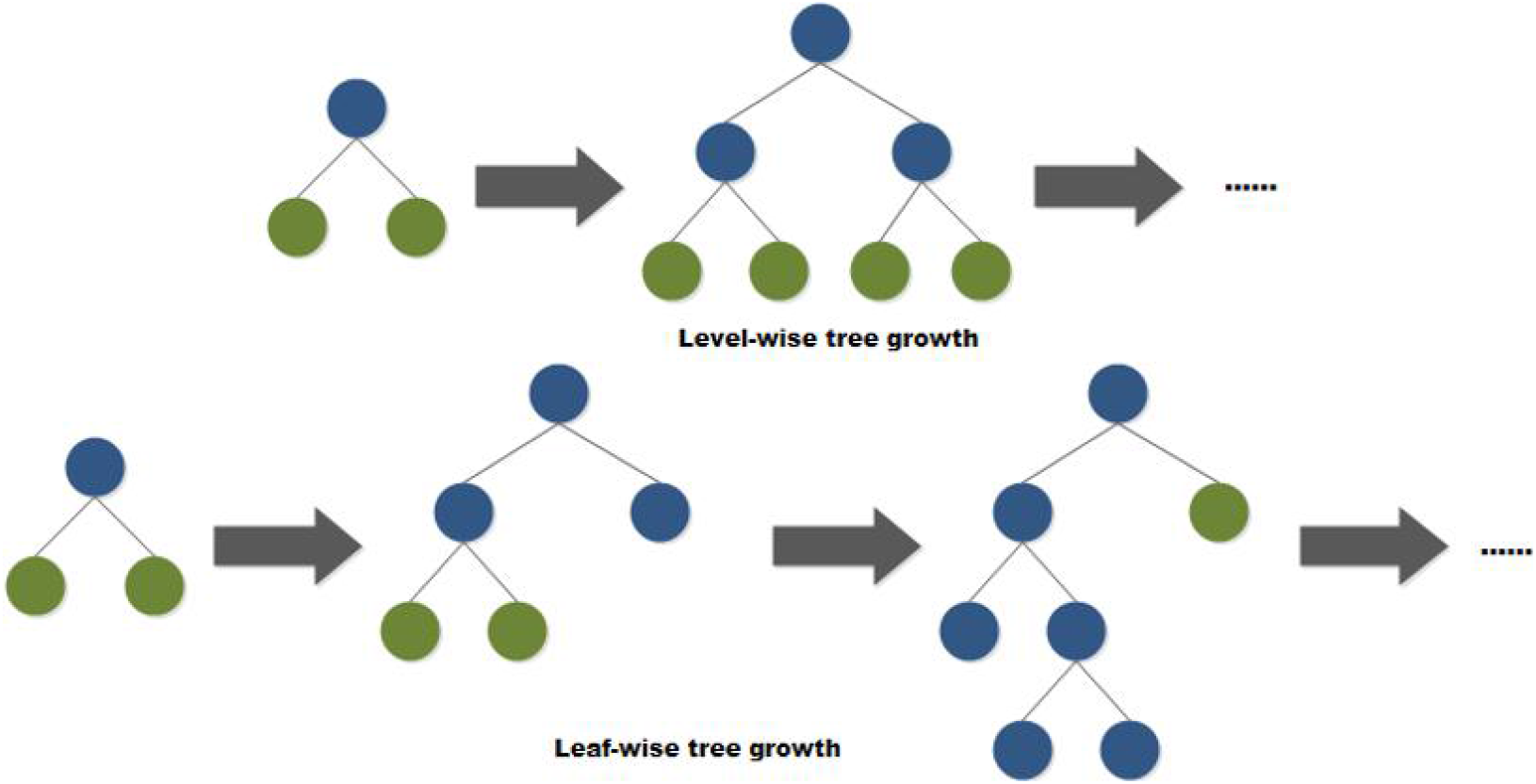
Comparison of Leaf-wise and Level-wise growth strategies.

##### 2.3.2.2 Extreme Gradient Boosting (XGBoost)

XGBoost is one of the boosting algorithms. It employs the sum of the predicted value of each tree in the K samples (the total number of trees is K) (that is, the sum of the scores of the corresponding leaf nodes of each tree) as the prediction. A new function *f* is added to the prediction in each iteration to minimize the objective function. At present, as a novel ensemble learning algorithm, XGBoost presents great results and is widely used in disease detection and other fields [24], but there is no report about its application on allergenicity prediction of food proteins.

##### 2.3.2.3 Random Forest (RF)

Random forest is a typical model of Bagging ensemble learning. It combines multiple weak classifiers, and adopts voting methods to make the final decision, therefore having higher accuracy and generalization. Random forest has been used in allergen prediction research and is the representative of traditional ensemble learning in this field [25].

#### 2.3.2. Previous Machine Learning Models

In order to compare the performance of the novel deep learning model and ensemble learning proposed in this paper, we adopted the three baseline machine learning algorithms (SVM, K-NN and Naive Bayesian (NB), which are often employed in previous similar studies.) [10,15,26]. SVM is a supervised learning algorithm that solves two or multiple classification problems. After introducing the kernel method, it can also be used to solve nonlinear problems. In this work, SVM with non-linear kernel was adopted. The principle of K-NN is relatively simple. The classifier calculates the distance between the feature values of the training data and new data, then selects K (K≥1) closest neighbors for classification or regression. NB performs well on small-scale data. It is usually applied in multi-classification tasks because it is suitable for incremental training and has low sensitivity.

### 2.4 Performance Evaluation of Models

In this study, accuracy (Acc), recall, precision (Prec), F1 score (The definition of these indicators are shown as follows) and area under the Receiver operating characteristic curve (ROC and AUC) were selected to evaluate the performance of the model. It should be noted that the classification threshold of the above indicators was uniformly set to 0.5.

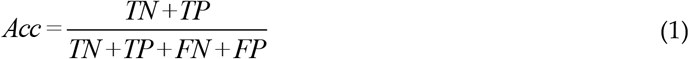

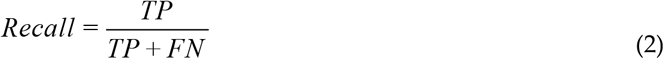

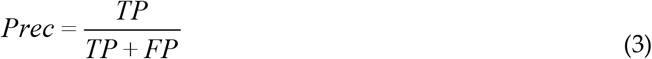

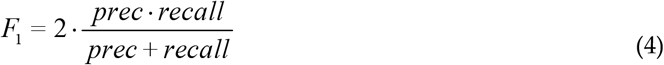

Where TN is the true negative number, TP is the true positive number, FN is the false negative number, and FP is the false negative number.

## 3. Results

### 3.1 Experimental Set Up

The ensemble and baseline models calculations was deployed in Windows 10 system, which was configured with CPU Intel Core I7-6700HQ, 3.5 GHz, 4 GB memory. Additionally, the related experiment of the deep model was performed on another equipment with better capability and the training process was powered by NVIDIA® Tesla T4 GPU, accelerated by CUDA. NVIDIA T4 is a universal deep learning accelerator which is widely used in distributed computing environments. The programming language used was Python 3.0 and Pytorch was chosen for the deep learning framework. In this study, each model was trained separately (BERT has been pre-trained), and the GridSearchCV interface in the scikit-learn third-party library was adopted for parameter optimization. 5-fold cross-validation was used for verification: the training set and the test set were randomly allocated at a ratio of 8:2 and repeated 5 times, and various evaluation indicators were recorded during the training. In order to reflect the performance of the model in real situations, we have calculated the mean value of each indicator 95.00% confidence interval (CI) for each model.

### 3.2 Performance of Deep Learning Models

By connecting left-to-right and right-to-left texts, a pre-processed deep two-way expression model was designed. After parameter optimization, the key parameters of the model were set as attention_probs_dropout_prob: 0.0, hidden_act: gelu, hidden_dropout_prob: 0.0, hidden_size: 1024, initializer_range: 0.02, intermediate_size: 4096, max_position_embeddings: 40000, num_attention_heads: 16, num_hidden_layers: 30, type_vocab_size: 2, and vocab_size: 30. The 5-fold cross-validation results of the deep learning model are shown in Table **1**. It can be found that the accuracy of the model reached 0.9310, the F1 score was 0.9344, which showed great generalization ability. Furthermore, the ROC curve of BERT and the corresponding AUC value are shown in Figure **5**. Its AUC reached 0.9578, showing the outstanding performance of our proposed method.

**Table 1.** BERT’s performance in the task of predicting food allergens.

| Model | Acc | Recall | Prec | F1 |
| --- | --- | --- | --- | --- |
| BERT | 0.9310±0.0145 | 0.9419±0.0163 | 0.9262±0.0203 | 0.9344±0.0141 |

**Figure 5.**
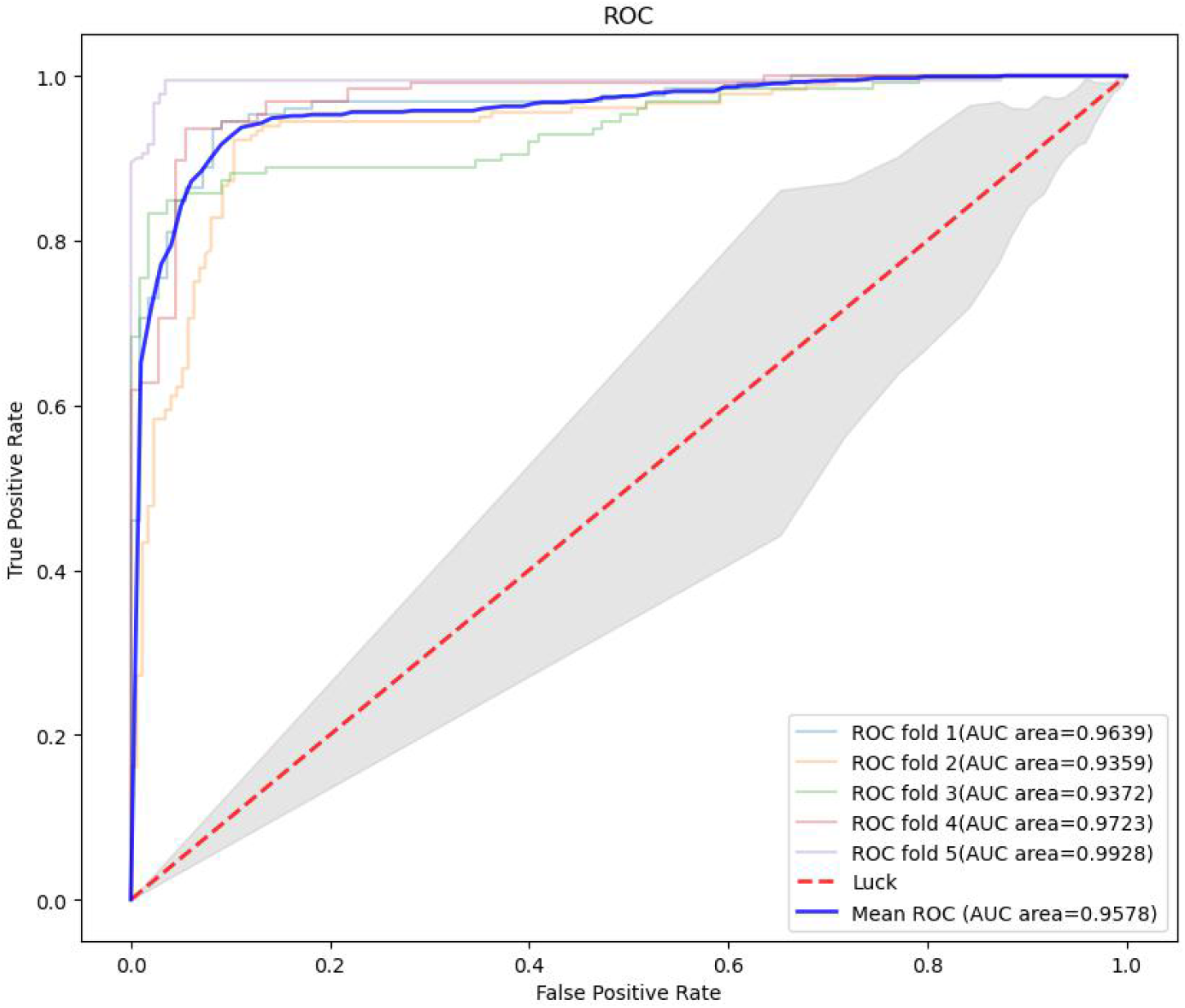
ROC curve and corresponding AUC value of BERT model.

### 3.3 Performance of Ensemble Learning Models

Training and verification for the three ensemble learning models mentioned above were conducted in the experiment, and each model has been optimized to a greater extent after adopting the parameter adjustment methods proposed above. Table **2** shows the key parameters of the ensemble models.

**Table 2.** The key parameters optimization results of the ensemble learning models.

| Model | Key Parameters Names and Corresponding Values |
| --- | --- |
| LightGBM | n_estimators=400, learning_rate=0.1, max_depth=5, num_leaves=32 |
| XGBoost | learning_rate=0.0001, n_estimators=1000, max_depth=5, subsample=0.8, seed=27 |
| RF | n_estimators=60, max_depth=13, min_samples_split=120, min_samples_leaf=20, max_features=7 |

The cross-validation results are shown in Table **3**. It can be seen that LightGBM and XGBoost performed best as novel ensemble algorithms. The average accuracy and F1 score of the two models were 0.8686, 0.8186 and 0.8684, 0.7981 respectively. RF model performed worse than the two. As a representative of Bagging ensemble models, the average accuracy and F1 score were only 0.7797 and 0.7720.

**Table 3.** Performance of the ensemble learning models in the task of predicting food allergens.

| Model | Acc | Recall | Prec | F1 |
| --- | --- | --- | --- | --- |
| LightGBM | 0.8686±0.0132 | 0.8793±0.0250 | 0.8571±0.0388 | 0.8684±0.0098 |
| XGBoost | 0.8186±0.0248 | 0.7778±0.0316 | 0.8426±0.0491 | 0.7981±0.0235 |
| RF | 0.7797±0.0370 | 0.7586±0.0449 | 0.7857±0.0544 | 0.7720±0.0182 |

In addition, the ROC curves and the corresponding AUC values of the models are shown in Figure **6**. LightGBM had the highest AUC value (0.9105), so its generalization ability was the best. The second was XGBoost (0.8803). It can be seen from the ROC curves and the corresponding AUC values that there were still some differences between the XGBoost and the LightGBM. The AUC of RF was 0.8542, which was quite different from LightGBM and XGBoost.

**Figure 6.**
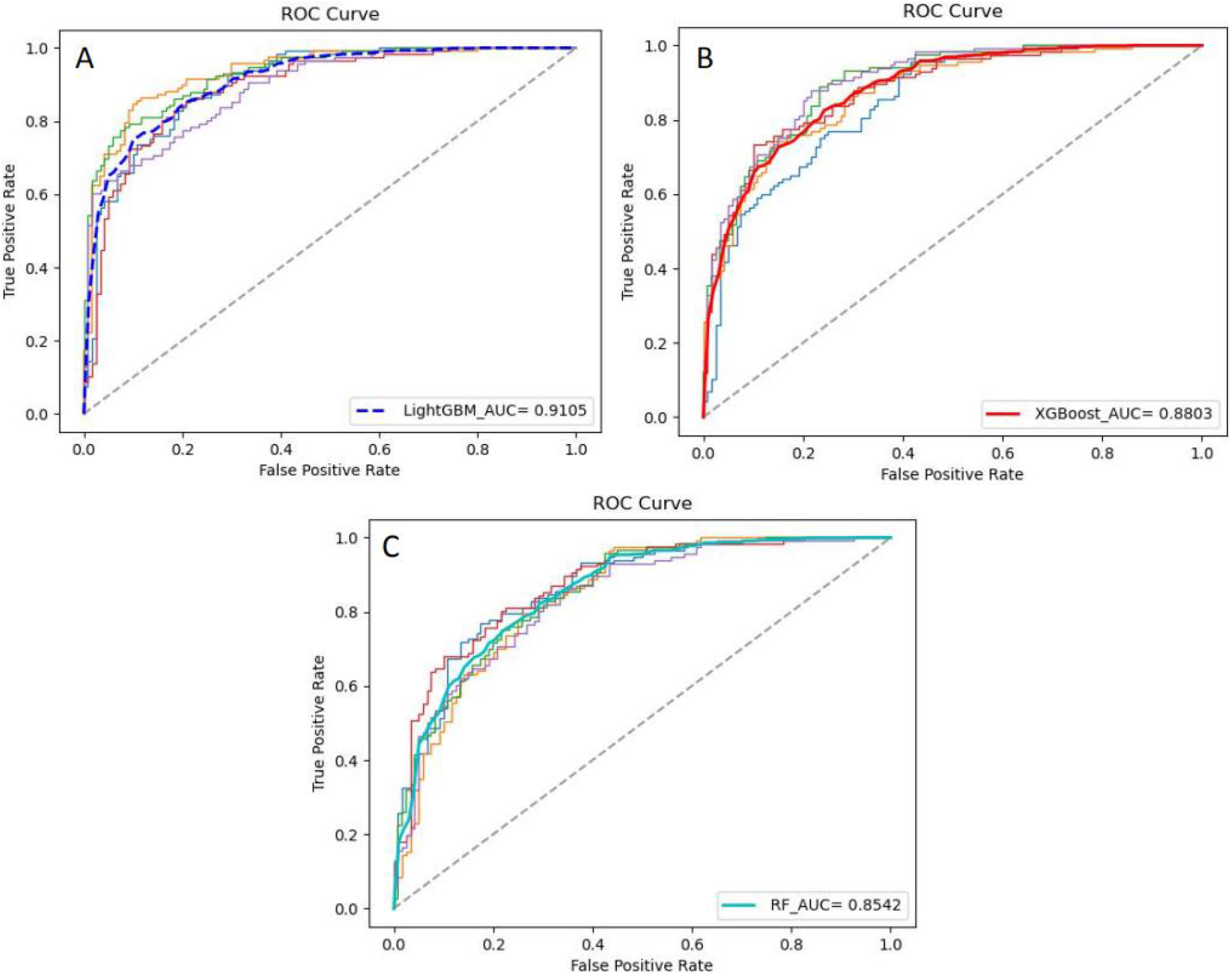
ROC curves and corresponding AUC values of the ensemble models.

### 3.4 Performance of Previous Machine Learning Models

As baselines for novel deep learning and ensemble learning models, the previously widely used allergen identification machine learning models (SVM, K-NN, NB) were also introduced in the experiment for comparison. This study extracted the pseudo-amino acid composition characteristics of the protein sequence, then input them into the classifier for training and optimization. The results of some parameters optimization are shown in Table **4**. The 5-fold cross-validation results are shown in Table **5**, and the ROC curve of each model and the corresponding AUC value are shown in Figure **7**.

**Table 4.** The key parameters optimization results of the previous machine learning models.

| Model | Key Parameters Names and Corresponding Values |
| --- | --- |
| SVM | C=1.0, kernel='rbf', gamma=0.01 |
| K-NN | n_neighbors=5, n_jobs=1 |
| NB | alpha=0.9 |

**Table 5.** Performance of previous machine learning models in the task of predicting food allergens.

| Model | Acc | Recall | Prec | F1 |
| --- | --- | --- | --- | --- |
| SVM | 0.7418±0.0420 | 0.7032±0.0443 | 0.7591±0.0524 | 0.7303±0.0389 |
| K-NN | 0.7722±0.0234 | 0.7436±0.0385 | 0.7838±0.0410 | 0.7630±0.0132 |
| NB | 0.7203±0.0375 | 0.6293±0.0466 | 0.7604±0.0517 | 0.6891±0.0221 |

**Figure 7.**
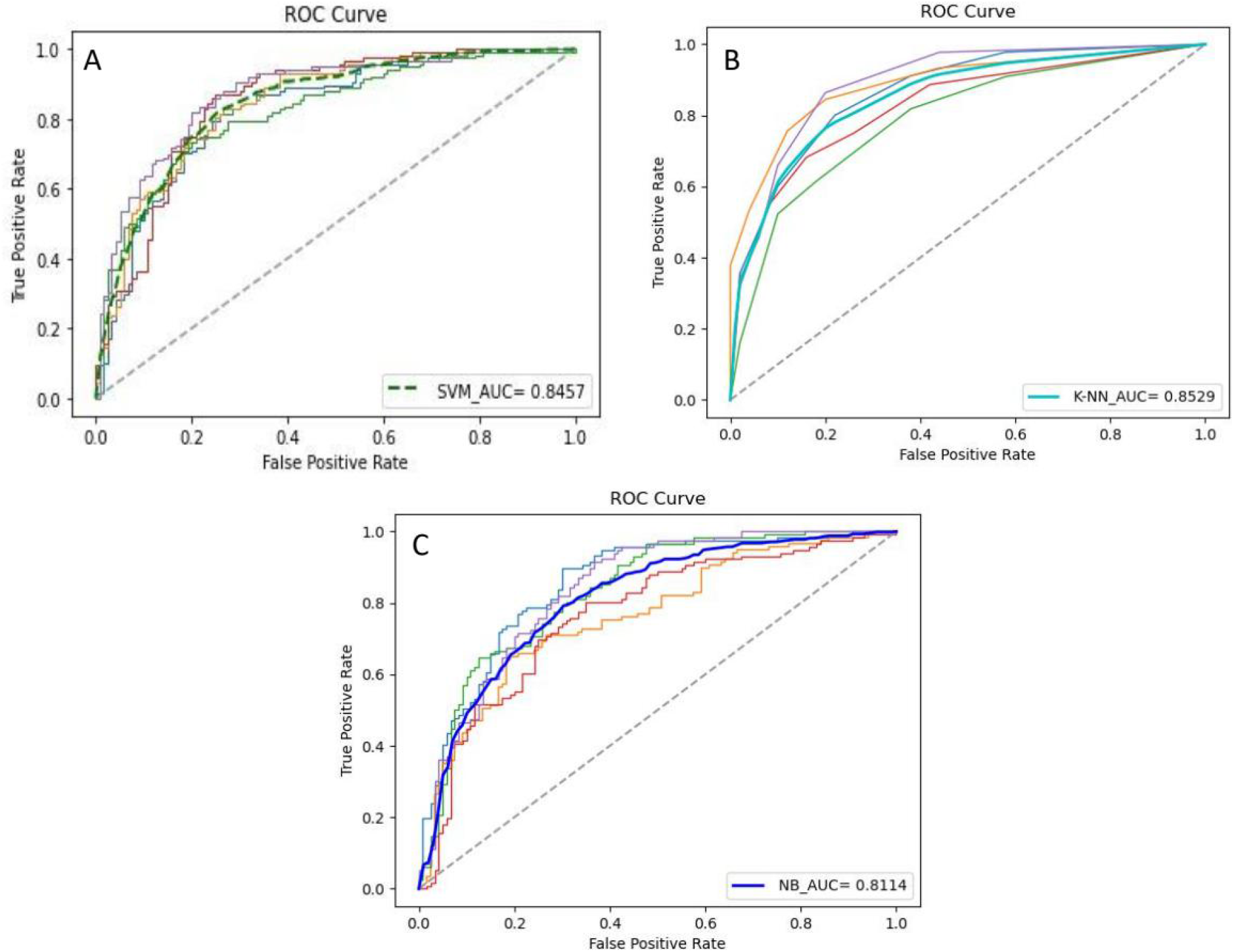
ROC curves and corresponding AUC values of the previous models

Compared with deep learning and ensemble learning models, the performance of the baseline algorithms was generally more inferior. Among them, the SVM achieved an accuracy of 0.7418 with an F1 score of 0.7303, and its AUC was 0.8457, which cannot make relatively accurate predictions for whether a test sequence is allergenic or not. In contrast, K-NN performed the best accuracy, perhaps due to the architecture of the model itself.

## 4. Discussion

In order to break through the bottleneck of low accuracy encountered by traditional allergen prediction methods, this work designed a deep learning model with novel self-attention transformer structure and improved tree ensemble models to predict which was superior to the machine learning methods employed in previous similar works for the allergenicity of food proteins. The work provided new ideas for future food allergen screening. As far as we know, this is the first reported work to introduce the BERT deep model, LightGBM and XGBoost ensemble models into the food allergen prediction task. In this section, we will compare and analyze the characteristics of the proposed models and discuss their application scenes, which will definitely facilitate the future model selection.

In the deep learning model of BERT, the advantage of introducing self-attention is that it can connect two long-term dependent features in the sequence. This may require more time to accumulate and react for the recurrent neural network (RNN) structure, so the self-attention mechanism can improve the parallelism of the network. The input of this research is protein sequences of different lengths. Self-attention can ignore the distance between amino acids and directly calculate their dependence relationship. It can help learn the internal structure of protein sequences well, which is better than traditional natural language processing algorithms and more efficient. Meanwhile, BERT model employed in this work has been pre-trained, and a large number of operations done in the downstream tasks of natural language processing are transferred to the pre-trained word vector. This not only improves the efficiency of the allergen sequence recognition, but also bestows it more powerful generalization ability. The architecture of BERT is based on multi-layer two-way conversion and decoding, where “two-way” means that when the model is processing a certain word (amino acid), it can use both of the previous word (amino acid) and the following word (amino acid) at the same time, which is different from traditional RNNs. The above advantages all highlight the great potential of BERT to accurately predict food allergens. In this study, BERT’s AUC reached 0.9578, which was better than all ensemble learning models, the best of which was 0.9105 and previously reported machine learning models, the best of which was 0.8529. The high AUC value shows its powerful predictive ability. In terms of recognition accuracy, BERT was 0.9310, which was also obviously excellent, better than LightGBM (0.8686) and XGBoost (0.8186). This benefits from the unique advantages of the transformer architecture, which surpasses the boosting ensemble models in the task of food allergen prediction. However, it cannot be ignored that pre-training requires a large amount of various types of protein sequences, which leads to a high cost of transfer learning. It must be emphasized that the BERT model has a large number of hyperparameters and requires a long training time (the training time was about 325 minutes in this study), which also puts forward strict requirements on computing equipment.

The novel ensemble learning models also performed well in the task of food allergens identification. For example, LightGBM is a novel GBDT algorithm framework that has many advantages. One is GOSS, the algorithm does not adopt the sample points to calculate the gradient, but samples the samples to calculate the gradient. The second is EFB, which means that certain features are bundled together to reduce the dimensionality of the features. In addition, using the Leaf-wise strategy for iteration can reduce errors as much as possible and get better accuracy. Based on the above characteristics, LightGBM needs a shorter training time and has better learning effect than traditional machine learning algorithms for food allergen prediction. In this research, the average prediction accuracy of the model was 0.8686, the F1 score was 0.8684, and the AUC reached 0.9105, which showed that it has the ability to accurately predict the allergenicity of a test sequence under small-scale training. Additionally, as a novel ensemble learning model, XGBoost has been widely used in many fields. The study found that it has a relative excellent performance in food allergen prediction tasks through extensive experiment. As for RF, it has a relatively large gap between its performance and the former two. Compared with the BERT deep learning model, although the performance of the ensemble learning models was not as good as the former, but the algorithms represented by LightGBM and XGBoost didn’t require pre-training, and the training time lasted shorter (the training took about 1-2 minutes in this paper). This means that they can complete the task of screening food allergens on portable devices and obtain considerable results.

Table **6** compares the characteristics of deep model, ensemble models and the traditional models more clearly, including the prediction effect, time-consuming (5-fold cross-validation) and corresponding computing equipment. It is undeniable that the BERT model with high training cost is more suitable for large-scale and high-standard food allergen screening, and the boosting models proposed is more suitable for rapid operation on simple equipment.

**Table 6.** Comparison of different types of models in this work.

| Model Type | Model Name | Prediction Accuracy (%) | Time-consuming | Computing Equipment |
| --- | --- | --- | --- | --- |
| Deep Learning | BERT | 93.10 | About 19 500s | NVIDIA® Tesla T4 GPU, accelerated by CUDA |
| Ensemble Learning | LightGBM | 86.86 | About 100s | CPU Intel Core I7-6700HQ, 3.5 GHz, 4 GB memory |
|  | XGBoost | 81.86 | About 125s |  |
|  | RF | 77.97 | About 90s |  |
| Previous Machine Learning | SVM | 74.18 | About 75s |  |
|  | K-NN | 77.22 | About 70s |  |
|  | NB | 72.03 | About 60s |  |

In previous similar studies, AllerHunter [27] employed a self-designed coding scheme and SVM algorithm as a classifier to predict allergens and achieved good results. The highest AUC value reached 0.928, which was lower than the AUC (0.9578) of the BERT deep model. [11] used the PseAAC encoding method and selected the SVM classifier to predict the allergenicity of allergen proteins with the highest AUC value, which was lower than the novel machine learning algorithm proposed in this paper. AllerTop [28] and AllerTop.v2 [29] received more approval for proposing convenient online servers for allergen screening with optimal algorithm K-NN. After training and optimization (5-fold cross-validation), the screening accuracy was 0.8530, which was lower than the deep learning model we proposed. Furthermore, researchers have also utilized the descriptor fingerprint method to classify allergens, achieving an identification accuracy of 0.8800 in a large-scale dataset. Based on this, they developed an online service system AllergenFP [14]. The deep model BERT employed in the study and the ensemble learning models represented by LightGBM and XGBoost further improved the performance of allergen prediction. In a relatively small dataset, it still achieved the highest AUC value of 0.9578 and the highest accuracy of 0.9310.

But it is undeniable that certain limitations still exist in this experiment. For example, since we focused on the development of rapid prediction methods for the allergenicity of food allergen, only food allergen sequences were considered in the establishment of the dataset, and the scale was small than the overall allergen. It should be emphasized that strict allergenicity prediction studies need to be verified by in vitro wet experiments (such as ELISA, etc.), which will be further improved in the future.

## 5. Conclusion

This work proposed to adopt the pre-training BERT deep learning model and novel ensemble learning models represented by LightGBM and XGBoost to predict the allergenicity of food proteins. Extensive experiments results in excellent results. They were superior to the previous studies. In the results, the AUC value of BERT (performed best) reached 0.9578, and the accuracy reached 0.9310. The experiments has been conducted to compare and analyze the characteristics of the different models and provides a guidance for the applicable scenarios. So as far as we know, this work is the first reported study of using the above method to identify the allergenicity of food proteins, which will provide an inspiration for food allergens prediction in the future.

## Author Contributions

Conceptualization, L.W. and D.N.; methodology, L.W. and X.Z.; software, L.W.; validation, D.N. and X.W.; formal analysis, D.N.; data curation, X.W.; writing—original draft preparation, L.W., H.C., and M.H.; writing—review and editing, L.W. and X.Z.; visualization, X.W. and X.Z.; project administration, H.C.; funding acquisition, H.C. All authors have read and agreed to the published version of the manuscript.

## Funding

This work was funded by the National Natural Science Foundation of China (Grant No. 81773435)

## Institutional Review Board Statement

Not applicable.

## Informed Consent Statement

Not applicable.

## Data Availability Statement

The data presented in this study are available on request from the corresponding author.

## Conflicts of Interest

The author reports no conflict of interest in this work.

## Notes

### Competing Interest Statement

The authors have declared no competing interest.

